# Disruption of flour beetle microbiota limits experimentally evolved immune priming response, but not pathogen resistance

**DOI:** 10.1101/2022.07.11.499528

**Authors:** Arun Prakash, Deepa Agashe, Imroze Khan

## Abstract

Host-associated microbiota play a fundamental role in the training and induction of different forms of immunity, including inducible as well as constitutive components. However, direct experiments analysing the relative importance of microbiota during evolution of different immune functions are missing. We addressed this gap by using experimentally evolved lines of *Tribolium castaneum* that either produced inducible immune memory-like responses (immune priming) or constitutively expressed basal resistance (without priming), as mutually exclusive strategies against *Bacillus thuringiensis* infection. We disrupted the microbial communities in these evolved lines and estimated the impact on the beetle’s ability to mount a priming response vs basal resistance. Populations that had evolved immune priming lost the ability to mount a priming response upon microbiota disruption. Microbiota manipulation also caused a drastic reduction in their reproductive output and post-infection longevity. In contrast, in pathogen-resistant beetles, microbiota manipulation did not affect post-infection survival or reproduction. The divergent evolution of immune responses across beetle lineages was thus associated with divergent reliance on the microbiome. Whether the latter is a direct outcome of differential pathogen exposure during selection or reflects evolved immune functions remains unclear. We hope that our results will motivate further experiments to understand the mechanistic basis of these complex evolutionary associations between microbiota, host immune strategies, and fitness outcomes.

## INTRODUCTION

Growing evidence reveals the critical role of microbiota in altering various aspects of host development, behaviour, and reproduction (Gould et al., 2018), as well as in training and induction of host immune responses (Zheng et al., 2020). In many species, including humans, the microbiota is required for successfully mounting different forms of immunity (e.g., innate vs adaptive) (Chudnovskiy et al., 2016; Karimi et al., 2009; Mazmanian et al., 2005; Muhammad et al., 2019), such that depletion or loss of microbial diversity can increase the vulnerability to pathogens (Dillon and Dillon, 2003; Engel and Moran, 2013). Gut bacteria can also influence tissues, cells and molecular pathways involved in gastrointestinal immunity, and changes in microbiome composition leads to overactive inflammatory responses causing bowel disorders (Kostic et al., 2014). Together, these results indicate an optimal association between host and microbiota forged over a long coevolutionary history (Lee and Mazmanian, 2010), to appropriately train and regulate immune responses (Belkaid and Hand, 2014; Thaiss et al., 2016). Recent studies also suggest a role for microbiota in inducing immune memory-like responses in insects (immune priming), whereby prior exposure to a low dose of infection improves survival against a lethal infection caused by the same pathogen later in life (Futo et al., 2015; Muhammad et al., 2019). Thus, host microbiomes appear to be generally important in shaping various forms of immunity across diverse taxa.

However, it is less clear whether microbiota are similarly important in shaping different host immune strategies. Host immune systems can evolve to new equilibrium states reflecting distinct immune strategies in response to different pathogen selection pressures (Mayer et al., 2016, Khan et al. 2017). Although experimental support is missing, host-associated microbiota, owing to their immunomodulatory role, might also exhibit correlated changes as host immune functions diverge (Zheng et al., 2020). However, although pathogen resistance is one of the major evolutionary advantages conferred by microbiota (McLaren and Callahan, 2020), there are no experiments to test whether or to what extent the role of microbiota varies across divergent forms of host immunity. We thus conducted a proof-of-principle study to analyse the impacts of microbiota in replicated experimental evolution lines of flour beetle *Tribolium castaneum* that separately evolved either constitutively expressed higher basal resistance, or inducible immune priming responses against their natural pathogen *Bacillus thuringiensis* (Bt) (Khan et al., 2017; Prakash et al., 2022). We disrupted the microbiome of these evolved lines to test whether their evolved immune strategies and fitness traits depended on the host-associated microbiome.

## MATERIALS & METHODS

We used previously described, replicate populations of *T. castaneum* that were infected in each generation with live Bt (strain DSM 2046), either with or without prior exposure to priming with heat-killed Bt cells to create distinct selection regimes (Khan et al., 2017; also see the supplementary information) as follows. (a) C populations: Control populations with no priming or infection; (b) PI populations: Priming with heat killed Bt, followed by infection with live Bt (PI); and (c) I populations: Mock priming (i.e., injected with insect Ringer), followed by infection with live Bt. Of the 4 original replicate populations per selection regime, in the present work, we analysed three replicates (total 9 populations). After 14 generations of continuous selection, we found that I populations only evolved priming responses, whereas PI beetles had higher basal resistance (Prakash et al., 2022) as mutually exclusive responses— i.e., evolved populations either showed priming or resistance, but never produced both the responses together. In this study, we used the same beetle lines after another round of selection (i.e., 15 generations), then removed pathogen selection for two additional generations to minimize maternal or other epigenetic effects. We then collected “standardized” eggs to obtain experimental beetles with minimum non-genetic parental effects, to analyse the impact of disrupting microbiota on the already evolved immune responses (i.e., priming vs basal resistance).

### Experimental manipulation of the beetle microbiome and subsequent assays

Previous work shows that the beetle microbiome is most likely acquired from the flour that the beetles inhabit and consume, and in which they also defecate and reproduce (Agarwal and Agashe, 2020). Beetles also derived significant fitness benefits from flour-acquired microbes, including higher fecundity and lifespan (Agarwal and Agashe, 2020). Thus, the easiest way to manipulate the beetle microbiome is to deplete the flour-associated microbial flora. We followed a previously published protocol in the lab (Agarwal and Agashe, 2020), where thin layers of wheat flour were exposed to UV radiation (UV -C ~254nm) in a laminar airflow for 2h. This treatment significantly alters flour microbiome with drastic depletion of the dominant bacterial taxa (also see Fig. S1 for reduction in CFUs on LB agar plates post UV-treatment). We then isolated single standardised eggs from each population in the wells of 96 well plates containing ~0.25 g of either UV-treated flour or normal wheat flour and reared them as virgins until adulthood. We did not track the sex of beetles in subsequent experiments (unless stated otherwise) because neither priming nor basal infection responses varied across sexes in our previous studies (Khan et al., 2017; Prakash et al., 2022). Below, we describe the assays performed with standardised beetles reared in normal vs UV-treated flour—

A. <u>Evolved priming vs basal infection response:</u> To prime and infect beetles, we used the septic injury method as described earlier (Khan et al., 2016; also see SI information). Briefly, 10-day old virgin I regime adults (24 beetles/priming treatment/ microbiota manipulation/ replicate population) were randomly assigned to one of the following treatments: beetles were either injected with insect Ringer solution (unprimed) or primed with heat-killed Bt cells adjusted to 10^11^ cells/100μl Ringer solution (primed). Six days later, we infected all beetles with live Bt (~10^10^ cells in 75 μl Ringer solution) and recorded their mortality for 14 days. We did not assay priming for C and PI beetles since they never showed a priming response in our earlier experiments (see the assay in generation 14, (Prakash et al., 2022)). Instead, we compared 16-day old C and PI unhandled beetles directly for survival after infection with live Bt, across microbiota manipulations (n=24 beetles/treatment/dietary resource/replicate populations). This is because evolved basal resistance of PI is an estimate relative to post-infection survival of control C beetles which did not evolve against Bt. We did not observe any mortality in sham-infected beetles. We analysed priming and basal infection response data using a mixed effects Cox model (implemented in R, Therneau, 2015) with replicate population as a random effect, specified as: (1) Priming ~ Priming treatment (i.e., unprimed vs primed) x microbiota manipulation (i.e., UV-treated vs normal wheat flour) + (1| replicate population)] (2) Basal infection response ~ Selection regime (i.e., C & PI) x microbiota manipulation + (1| replicate population). A significant interaction between priming treatment (or selection regime) and microbiota manipulation would indicate that the survival benefits of evolved priming (or evolved basal infection response) in I (or PI) populations vary significantly with disruption of microbiota. Further, to disentangle the changes in priming response of I beetles with vs without the microbiota manipulation, we analysed priming for each microbiota manipulation treatment separately, using a mixed effects Cox model specified as: Priming ~ Priming treatment + (1| replicate population), with priming treatment and replicate population as a fixed and random effect respectively.
B. <u>Lifespan after priming:</u> In a separate experiment, we collected virgin females reared in normal vs UV-sterilized wheat flour as described above (n= 12 females/treatment/microbiota manipulation/replicate population) to estimate the long-term survival benefits of priming (same dose as mentioned above) and basal infection response against a lower dose of infection adjusted to 10^6^ cells in 75μl Ringer solution. We observed beetle mortality every 5 days until 90 days when most of them were dead. We analysed lifespan data using model specifications as described above.
C. <u>Reproductive output:</u> Finally, we measured the impact of microbiota manipulation on reproductive fitness of evolved PI and I beetles. We first paired 10-day-old unhandled virgin males and females across selection regimes and microbiota manipulation. After two days of mating, we separated the females and allowed each to oviposit for 48h in 5g wheat flour (n=39-52/treatment/replicate population). After 4 weeks, we counted the total number of eggs laid per female as a proxy for reproductive fitness. At each step, beetles were either given access to UV-irradiated or untreated flour, according to their rearing condition. We analysed the data using a mixed effects Generalised Linear Model with Quasi-Poisson error, specified as: Reproduction ~ Selection regime (i.e., C, I, PI) x microbiota manipulation + (1| replicate population). To disentangle the changes in each selection regime, we also analysed them separately.

For each analysis, we could pool the data across replicate populations since population identity did not show any significant main impact or interactions as a fixed factor (P<0.05).

## RESULTS

### I. Disruption of microbiota causes the loss of immune priming response but not basal resistance

Here, we present results from data pooled across replicate populations of each selection regime, since we did not find a significant population effect (see Methods). Separate analyses and plots for each replicate population are shown in the supplementary materials. We first compared the priming response of I beetles (with an evolved priming response) reared in normal vs UV-treated flour. We found a significant interaction between priming treatment and microbiota manipulations (Table S1A). Priming improved beetle survival only in I populations reared in the standard diet (normal wheat flour with microbes), but not when they consumed UV-treated flour (Fig. 1A, S2A; Table S1B, S1C). Thus, there was a loss of evolved priming ability with disruption of the dietary source of microbiota.

**Figure 1.**
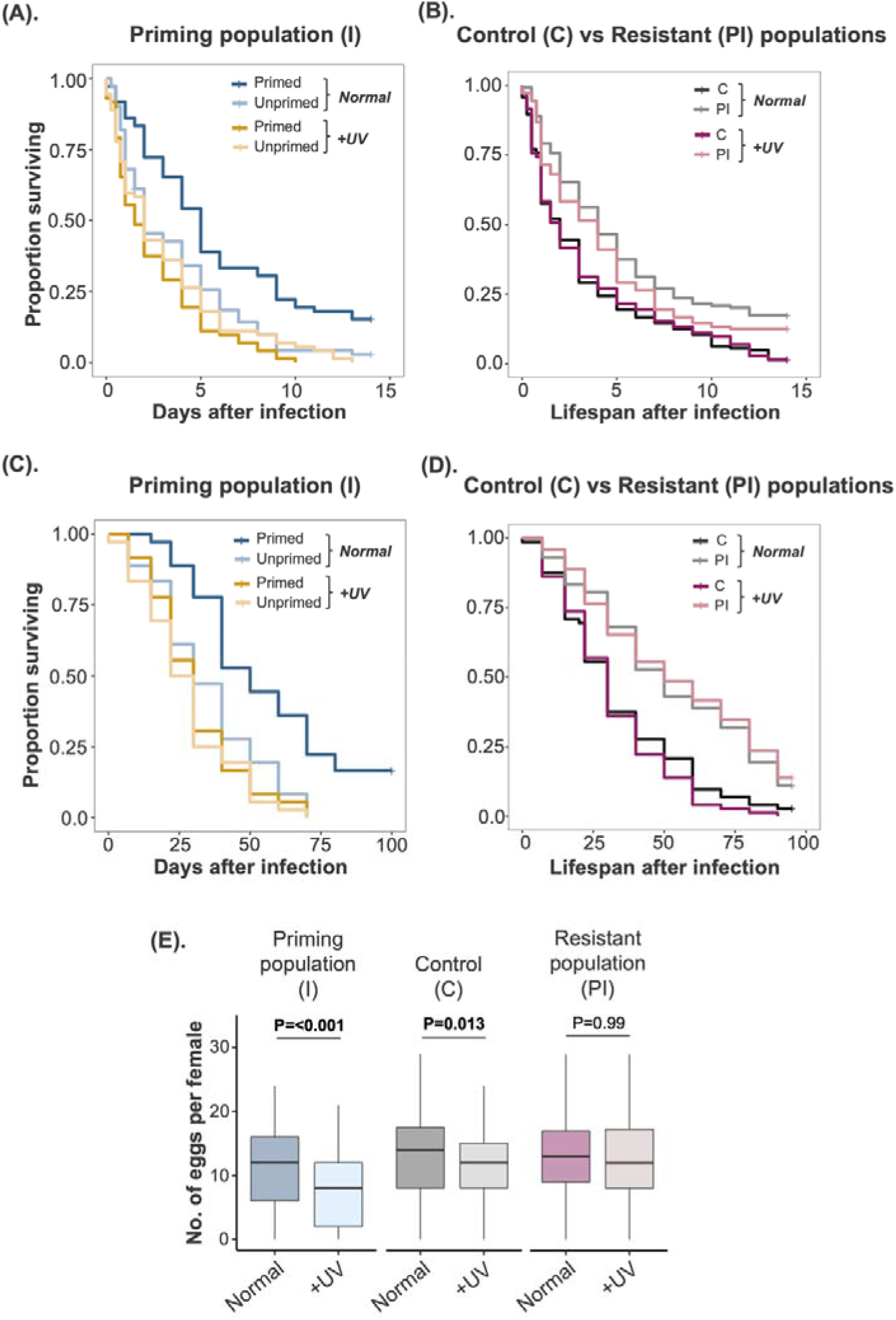
Effects of microbiota disruption by UV-irradiation of flour on (**A**) priming response (i.e., post-infection survival of primed vs unprimed individuals) of I beetles (with evolved priming); (**B**) basal resistance to Bt, i.e. post-infection survival of C (Control beetles) vs PI populations (with evolved higher basal resistance); (C) Lifespan of I beetles after priming and low dose of Bt; (D) Lifespan of C vs PI beetles after low dose of infection; (E) Reproductive fitness of naïve beetles from C, PI and I populations. In panel E, asterisks indicate significantly different groups. C= Control populations; I (or PI) = Replicate populations that evolved priming (or strong basal resistance); Normal= untreated wheat flour; +UV= UV-irradiated wheat flour; ns= not significant. Each panel represents pooled data across replicate populations (see Figs S2–4 for individual replicate populations). For panels A and B, n=24 beetles/priming and infection treatment/microbiota manipulation/replicate population; for panels C and D, n=12 females/priming and infection treatment/microbiota manipulation/replicate population; for panel E, n=39-52/microbiota manipulation/replicate population.

Subsequently, we compared C vs PI beetles to estimate the changes in basal resistance as a function of the microbiota manipulation. We found a significant main effect of the selection regime (as expected, PI beetles had higher, evolved basal resistance), but microbiota manipulation had no impact (Table S2). Further, the lack of a significant interaction between selection regime and microbiota manipulations indicated that the higher post-infection survival of PI beetles was not affected by their microbiota (Fig. 1B, S2B; Table S2).

These results corroborate another independent experiment where selected beetles were infected with a relatively lower dose of Bt and their lifespan was recorded until 90 days post-infection. Primed I beetles lived significantly longer when they were reared in the standard diet, but this benefit of priming disappeared when beetles were fed with UV-treated flour (Fig. 1C; S3A; Tables S3A, S3B, S3C). Hence, the longevity effects of priming also relied on the presence of microbiota. As expected, PI beetles lived longer than C beetles, regardless of the UV treatment of their diet (Fig. 1D, S3B; Table S4), suggesting that longevity effects of basal resistance do not depend on dietary microbes.

### II. Disrupting dietary microbes affects the reproductive potential of beetles with evolved priming, but not that of resistant beetles

Next, we analysed the effects of dietary microbe manipulations on reproductive output of beetle populations with divergent immune functions. Depletion of microbiota reduced reproductive output in both C and I females, but not in PI females (Fig. 1E, S4; Table S5A). Interestingly, the negative effect of microbiome disruption was more pronounced in I beetles, with a steeper decline of fitness relative to C beetles (Fig. 1E). We also note that the results for pooled data of C populations (Fig. 1E) differ from individual replicate populations (Fig. S4; Table S5B), which separately did not show a significant impact of microbiota manipulation. However, all the populations showed a consistent trend towards lower reproductive output of C beetles in UV treated flour (Fig. S4).

Interestingly, C beetles (which evolved in the absence of pathogen selection) most closely represent the ancestral condition. Thus, starting from a baseline negative effect of microbiome loss on reproduction, in I populations the effect became more pronounced, whereas in PI populations the effect of microbiota was lost. Thus, the reproductive effects of microbiota potentially co-evolved with pathogen selection and host immune strategies.

## DISCUSSION

In the past few decades, we have learnt that host-associated microbial communities can have major impacts on the host immune system and may have co-evolved with their hosts over evolutionary time (reviewed in Zheng et al., 2020). However, we lack direct evidence for the impact of microbiota on different components of the immune system when evolving under strong pathogen selection. This is possibly due to the lack of a suitable experimental system where evolutionary trajectories of different immune responses can be clearly distinguished. Previously, we reported a unique set of experimentally evolved beetle lines where pathogen-imposed selection led to the rapid, parallel and divergent evolution of either strong basal pathogen resistance (PI populations) or immune priming (I populations) as mutually exclusive responses (Khan et al. 2017; Prakash et al., 2022). Here, we showed that evolved basal resistance vs priming ability also have varied functional dependence on the host beetle microbiota. The disruption of microbiota led to a complete loss of the survival benefit of priming, whereas basal resistance to Bt infection remained unaffected. In beetles that evolved priming ability, depletion of microbiota also revoked the benefit of longer lifespan after priming and reduced their reproductive output; but this was not the case in resistant PI beetles. Impacts of microbiota as a function of evolved immune responses might thus extend to multiple fitness traits. Moreover, the absence of reproductive effects in PI beetles starkly contrasts the observation that in unselected control C beetles an intact microbiota was necessary to maintain reproductive output. PI beetles thus also gained independence from the reproductive fitness effects of microbiota during evolution of basal resistance against pathogens.

Why does the effect of microbiome differ across evolved immune strategies? We speculate that the effects may be determined by how the microbiome modulates specific immune pathways underlying priming or basal infection responses. For example, in flour beetles, prior priming improves post-infection survival by controlling pathogen growth (Khan et al., 2019), with the help of canonical resistance mechanisms such as increased phenoloxidase response (Ferro et al., 2019). However, if the disruption in microbiome composition interferes with the activation of bactericidal phenoloxidase response in primed I beetles, they may have a higher pathogen burden, thereby neutralizing the net beneficial effects of priming. Recent experiments with the moth *Plodia interpunctella* corroborate this hypothesis: removal of gut bacteria reduced phenoloxidase activity and concomitantly increased mortality after Bt infection (Orozco-Flores et al., 2017). Dietary microbes may also somehow modulate the priming effects of Bt cells introduced into the beetle haemolymph via septic injury in our experiments, with the exciting implication of cross-talk between the gut environment and priming responses produced in the haemolymph (Freitak et al., 2007; Kwong et al., 2013). In contrast, the evolved basal infection resistance of PI beetles might have been achieved by improving overall body condition (Prakash et al., 2022), which could have also increased their ability to withstand the effects of infection (.e., increased tolerance, Seal et al., 2021), without directly activating or involving the immune response. As a result, the immunomodulatory effects of microbiota might not be relevant for PI beetles anymore. Another possibility is that distinct sets of microbes may regulate the efficiency of evolved basal resistance vs priming, with the former being UV-resistant and the latter UV-sensitive.

Finally, we note that the potential divergence in microbiomes as well as beetle immune function may be unlinked, with each being driven independently by the specific selection regime. Whether this hypothesis is true, and if so, what is the direction of causality, remains to be determined. For instance, the beetle microbiome could be first rapidly altered by Bt infection (Li et al., 2020), due to infection-induced changes in host physiology. Since I vs. PI regimes involved differential exposure to Bt, the two regimes may have allowed for divergent changes in the microbiome. Eventually, the altered microbiomes could have facilitated the subsequent evolution of beetle immune function. Alternatively, beetle immune function may have diverged first across regimes (Cherif et al., 2008), changing the resident microbiomes later as a by-product. To distinguish between these alternatives, one would need to analyse the time course of change in host immune function as well as microbiomes during evolution. We hope that our results revealing the possibility of divergent impacts of microbiota across immune strategies will spur further work to test whether or to what extent these changes in the host immune function and microbiome are causally linked, and if so, through what mechanism.

## Acknowledgements

We thank Biswajit Shit, Devashish Kumar, Mrudula Sane, Pratibha Sanjenbam, Shivansh Singhal, Shyamsunder Buddh, and Srijan Seal for feedback on the manuscript. We thank Kunal Ankola, Pavan Thunga, Sunidhi Thakur and Shyamsunder Buddh for laboratory assistance.

## Author contributions

IK conceived the experiment; IK, AP and DA designed the experiment; AP performed the experiment; AP and IK analyzed the data. IK & DA wrote the manuscript with inputs from AP.

## SUPPLEMENTARY INFORMATION

### Supplementary methods

#### I. <u>Priming and infection protocol</u>

We used a strain of *Bacillus thuringiensis* (Bt - DSM 2046), isolated from a Mediterranean flour moth (Roth et al., 2009), as a model bacterial pathogen to prime and infect adult beetles (see Khan et al., 2017). To prime beetles, we pricked them between their head and thorax with a 0.1 mm insect pin (Fine Science Tools, CA) dipped in heat-killed bacterial slurry adjusted to 10^11^ cells/100μl Ringer solution, prepared from freshly grown overnight Bt culture at 30°C (optical density OD_600_ = 0.95). We used insect Ringer solution as mock priming (unprimed). The priming with heat-killed Bt cells can activate the immune response without imposing any direct cost of infection. Six days after priming, we infected both primed and unprimed individuals with a live bacterial culture adjusted to ~10^10^cells in 75 μl insect Ringer solution. To measure lifespan after mounting a priming response, we used a milder dose of live bacterial culture adjusted to ~10^6^ cells in 75 μl insect Ringer solution which does not cause any immediate mortality (within 7 days).

#### II. <u>Experimental evolution protocol (see Khan et al., 2017 for detailed protocol)</u>

Briefly, at every generation of experimental evolution, we primed 10-day-old virgin PI adults from each replicate population with heat-killed Bt, as described above. Simultaneously, we also mock-primed 10-day-old virgin adult C and I beetles with sterile insect Ringer solution. After six days, we challenged individuals from I and PI regimes with high dose of live Bt infection as described above, whereas C beetles were just pricked with sterile insect ringer solution (mock challenge). Hence, we had two infection regimes where populations were challenged with a high dose of Bt infection, with (PI populations) or without (I Populations) the opportunity of priming; and a control regime (C populations) where beetles were never exposed to Bt antigen. Following the priming and infection treatments, we combined 60 pairs of surviving males and females from each replicate population and allowed females to oviposit for 5 days to initiate the next generation. We repeated the same protocol for 15 generations, and then allowed two generations of relaxed selection (i.e., no pathogen exposure) before we commenced the assays described in this study.

## Supplementary figures

**Figure S1.**
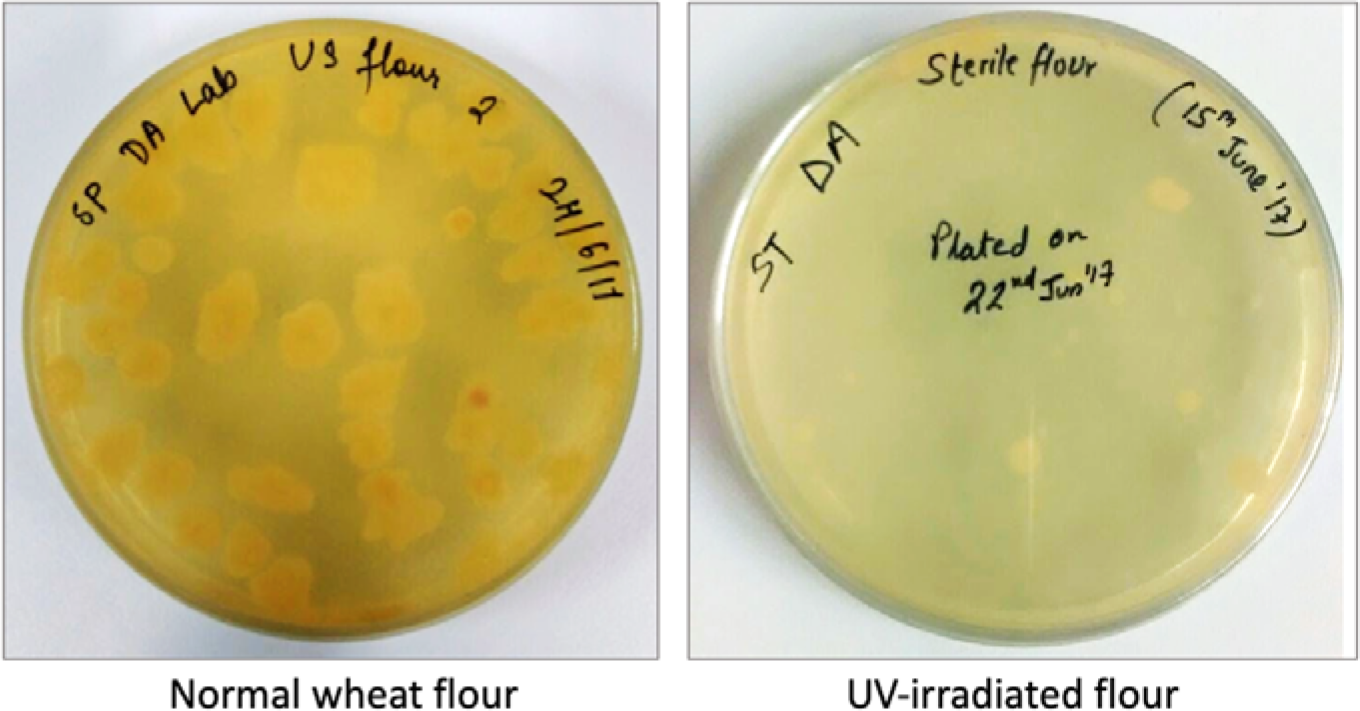
Representative LB agar plates to show the depletion of culturable microbiota after UV-irradiation of wheat flour

**Figure S2.**
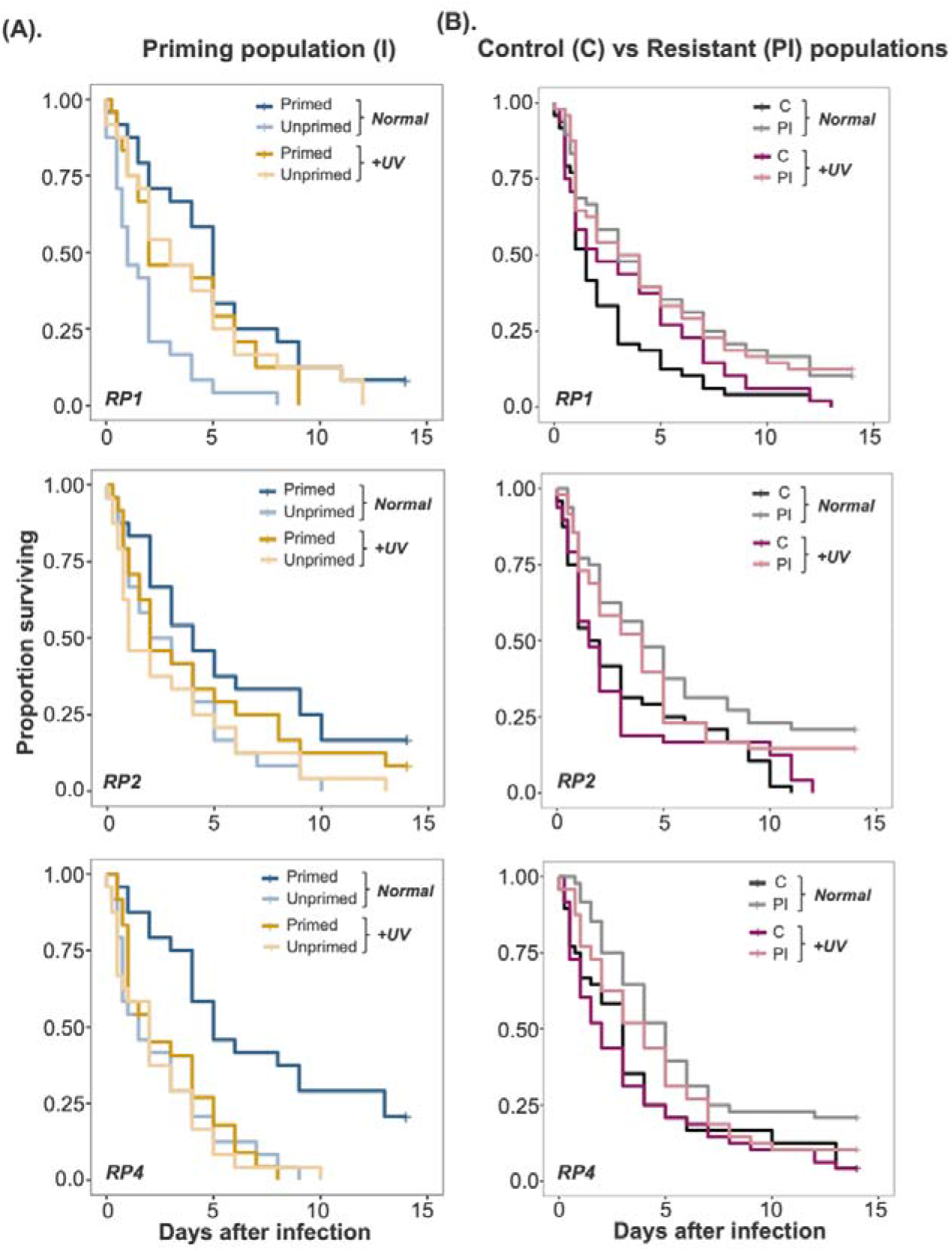
Effects of microbiota disruption by UV-irradiation on (A) priming response (measured as the difference between post-infection survival of primed vs unprimed individuals) of each beetle replicate populations that evolved priming (I1, 12, 14 replicate populations) (n=24 beetles/priming and infection treatment/microbiota manipulation/replicate population); (B) Post-infection survival of beetles from each replicate control population (C1, 2 & 4 populations) vs beetles that evolved strong basal resistance (PI1, 2, 4 populations) (n=24 beetles/infection treatment/ microbiota manipulation/replicate populations). C1 and PI1, C2 and PI2, and C3 and PI3 were handled together during the experiment. Normal= Normal wheat flour; +UV= UV-irradiated wheat flour.

**Figure S3:**
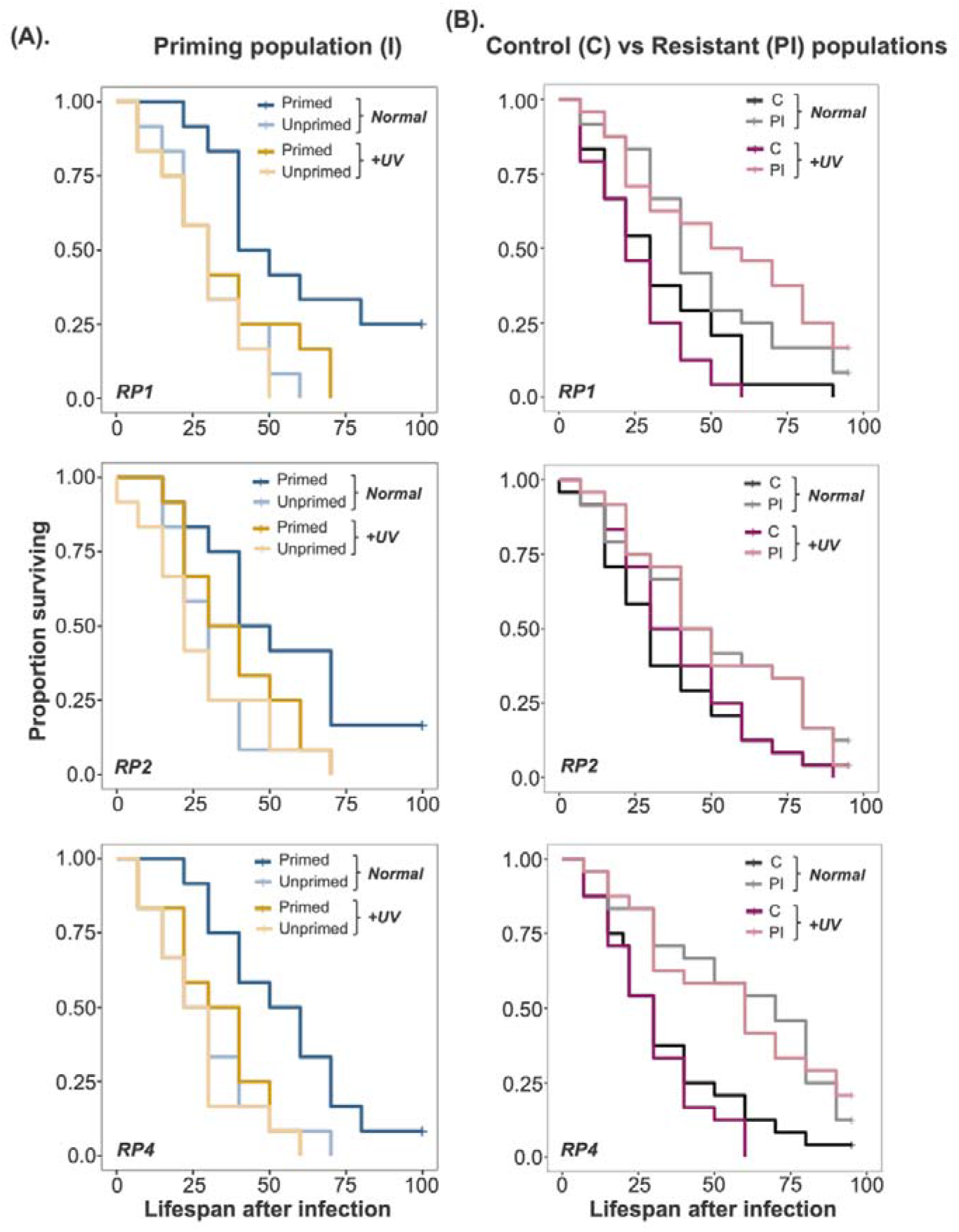
Effects of microbiota disruption by UV-irradiation on (A) lifespan after priming response (measured as the difference between post-infection lifespan of primed vs unprimed individuals) of each beetle replicate populations that evolved priming (I1,12, 14 populations) (n=24 beetles/priming and infection treatment/ microbiota manipulation/replicate population); (B) post-infection lifespan of each replicate populations of control beetles (C1, 2 & 4 populations) vs beetles that evolved strong basal resistance (PI1, 2, 4 populations) (n=24 beetles/infection treatment/ microbiota manipulation/replicate populations). C1 and PI1, C2 and PI2, and C3 and PI3 were handled together during the experiment. Normal= Normal wheat flour; +UV= UV-irradiated wheat flour.

**Figure S4:**
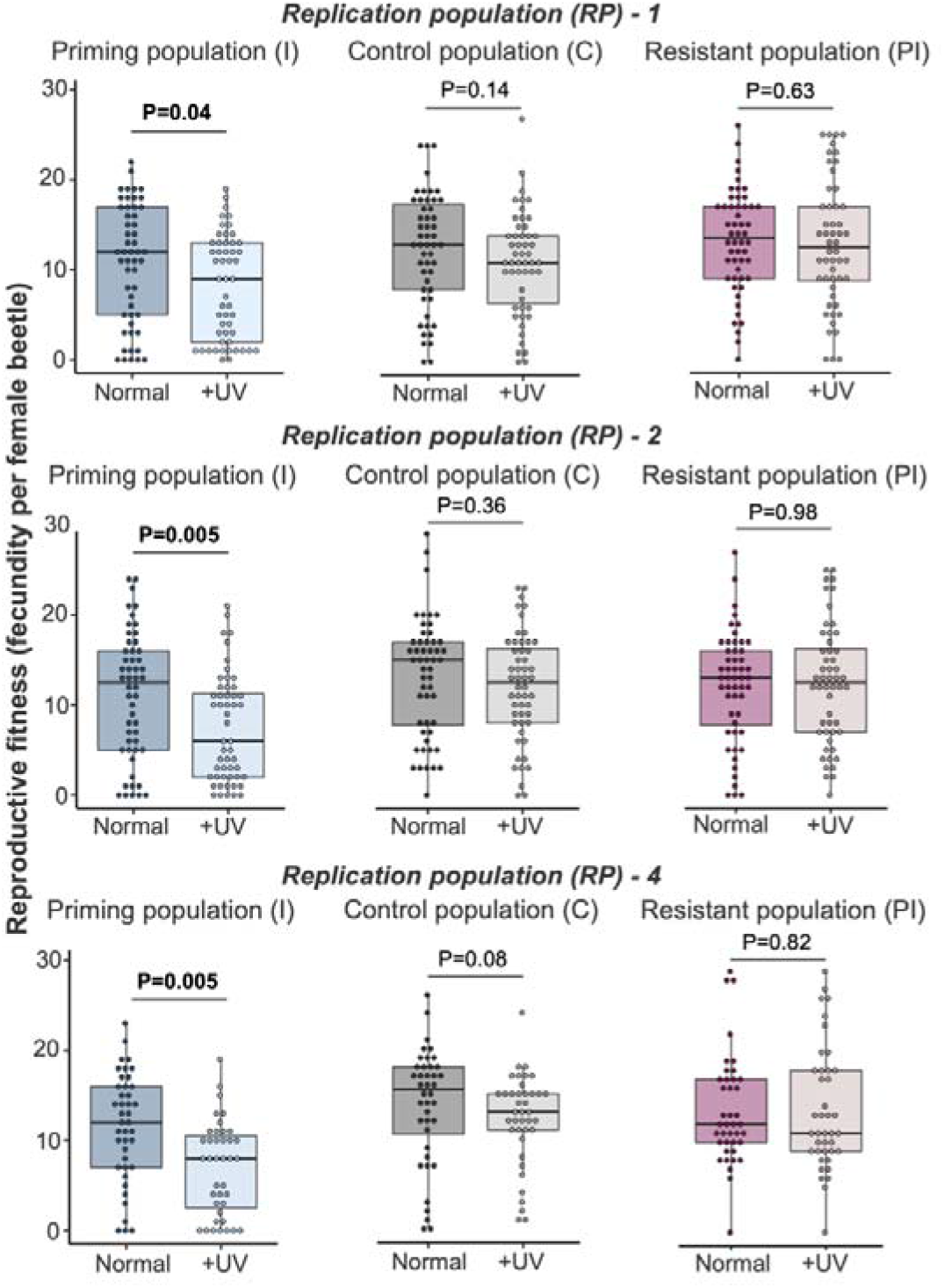
Effects of microbiota disruption by UV-irradiation on reproductive fitness of naïve beetles from each replicate populations of C, PI, and I populations (n=39-52 females/treatment/replicate population). Normal= Normal wheat flour; +UV= UV-irradiated wheat flour.

## Supplementary tables

**Table S1.**
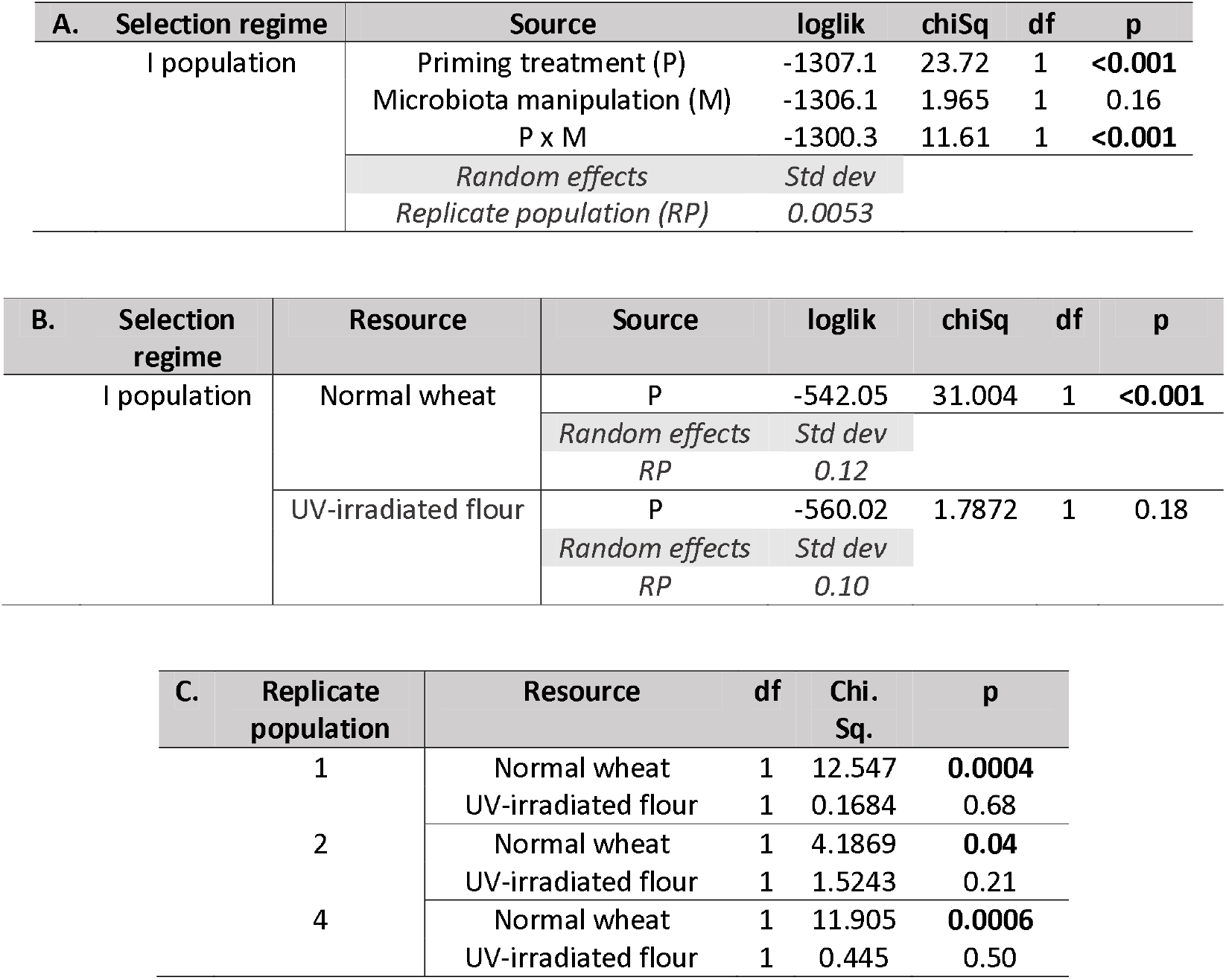
Summary of a mixed effects Cox model analysis to estimate the changes in priming response of I (experimentally evolved priming) population (A) as a function of microbiota disruption. We specified the model as: Priming response ~ Priming treatment (P) x microbiota manipulation (M) + (1| replicate population (RP))], with ‘P’ and ‘M’ as fixed effects, and RP as a random effect; (B) separately across microbiota manipulations (i.e., normal vs UV-irradiated flour). For each microbiota manipulation type, we specified the model as: Priming response ~ Priming treatment (P) + 1| replicate population (RP)], with ‘P’ as a fixed effect, and RP as a random effect; (C) Summary of a Cox proportional hazard analysis for priming response in each of the replicate | populations after disruption of dietary microbes.

**Table S2.**
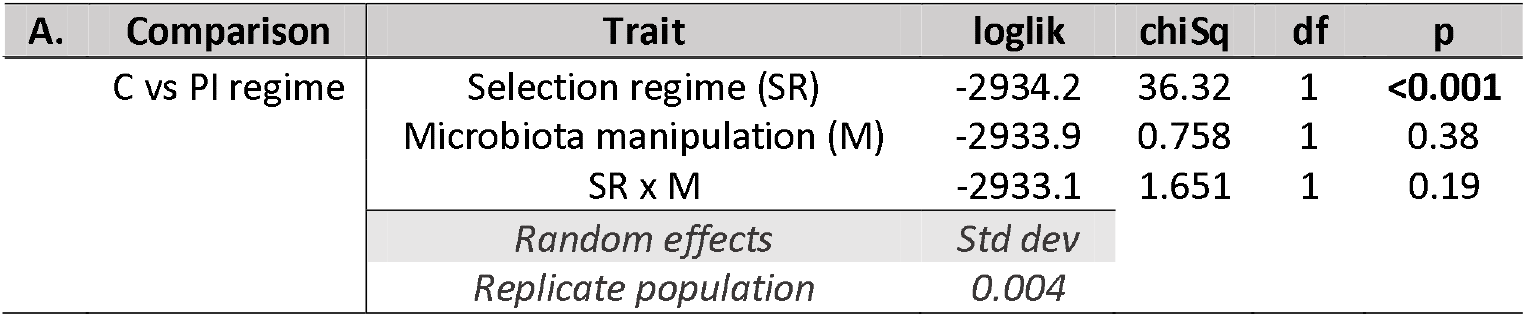
Summary of a mixed effects Cox model analysis on survival data of beetles from control (C) vs resistant (PI) populations as a function of microbiota manipulation. We specified the model as: Post-infection survival ~ Selection regime (SR) x microbiota manipulation (M) + (1| Replicate population (RP)), with ‘SR’ and ‘M’ as fixed effects, and RP as a random effect.

**Table S3.**
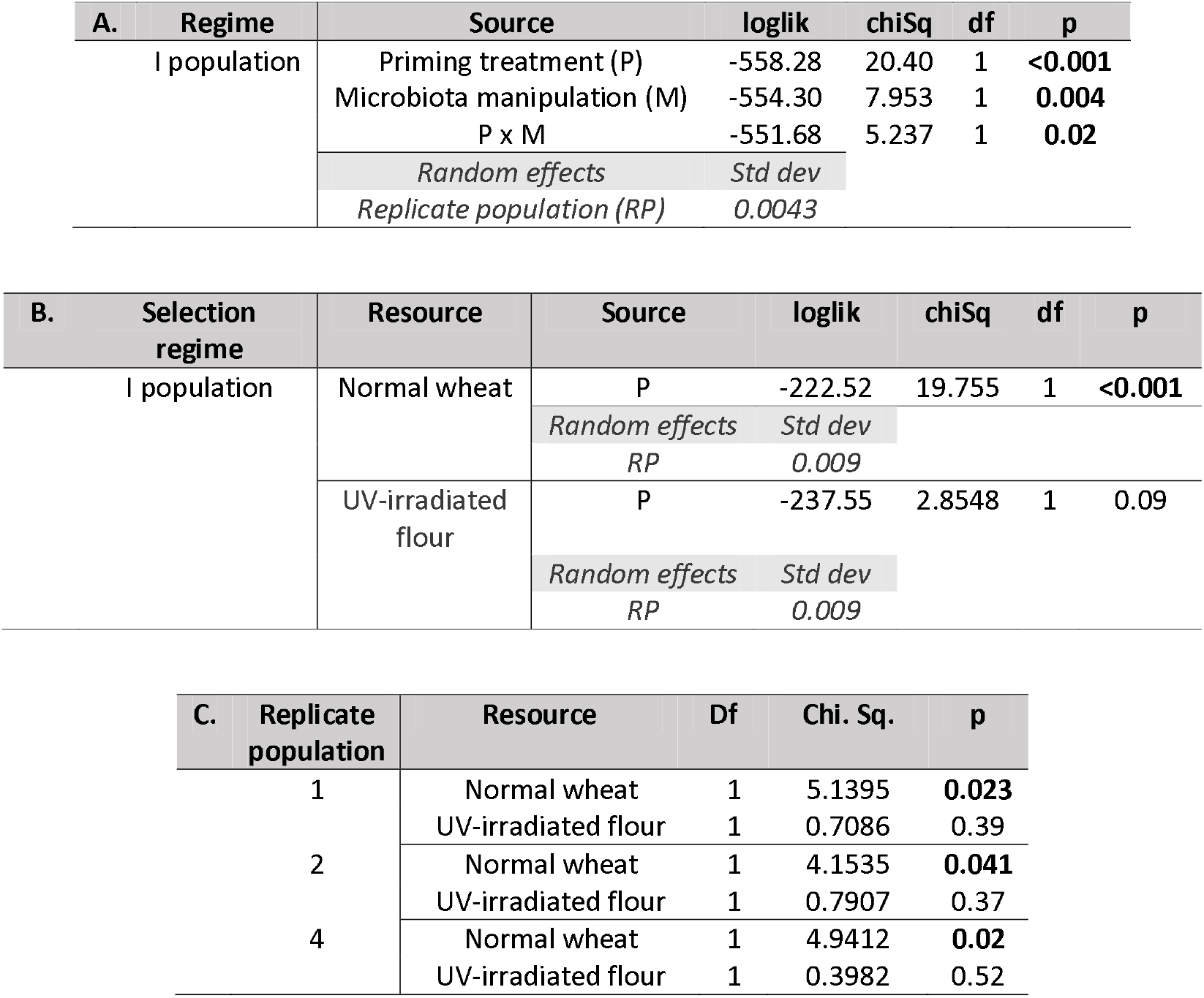
Summary of a mixed effects Cox model analysis to estimate the changes in lifespan of I beetles after priming response (A) as a function microbiota disruption. We specified the model as: Lifespan ~ Priming treatment (P) x Microbiota manipulation (M) + (1| Replicate population (RP))], with ‘P’ and ‘M’ as fixed effects, and RP as a random effect; (B) separately across microbiota manipulations (i.e., normal vs UV-irradiated flour). For each microbiota manipulation type, we specified the model as: Lifespan ~ Priming treatment (P) + 1| Replicate population (RP)], with ‘P’ as a fixed effect, and RP as a random effect; (**C**)Summary of a Cox proportional hazard analysis for lifespan after priming response in each of the replicate I populations after disruption of dietary microbes.

**Table S4.**
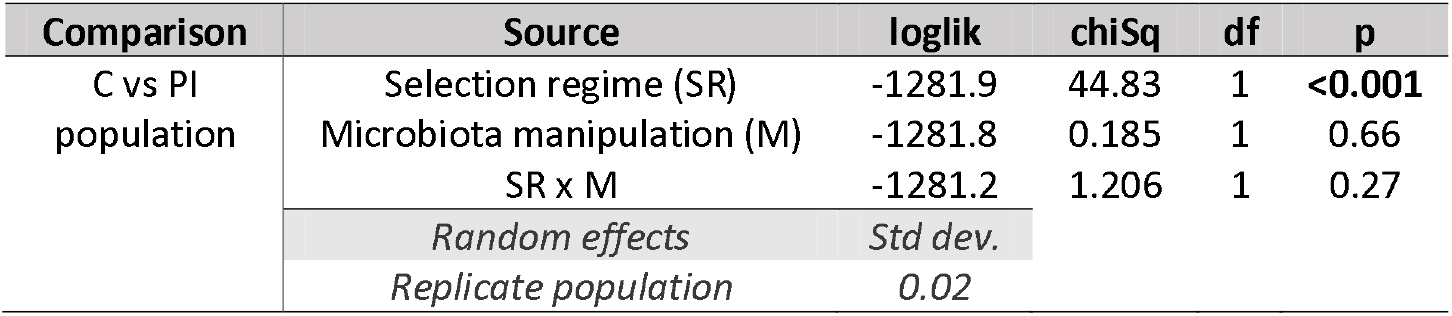
Summary of a mixed effects Cox model analysis on lifespan data of beetles from control (C) vs resistant (PI) populations after a mild infection dose, as a function of microbiota manipulation. We specified the model as: Lifespan ~ Selection regime (SR) x Microbiota manipulation (M) + (1| Replicate population (RP)), with ‘SR’ and ‘M’ as fixed effects, and RP as a random effect.

| Comparison | Source | loglik | chiSq | df | p |
| --- | --- | --- | --- | --- | --- |
| C vs PI population | Selection regime (SR) | -1281.9 | 44.83 | 1 | <0.001 |
|  | Microbiota manipulation (M) | -1281.8 | 0.185 | 1 | 0.66 |
|  | SR x M | -1281.2 | 1.206 | 1 | 0.27 |
|  | Random effects | Std dev. |  |  |  |
|  | Replicate population | 0.02 |  |  |  |

**Table S5:**
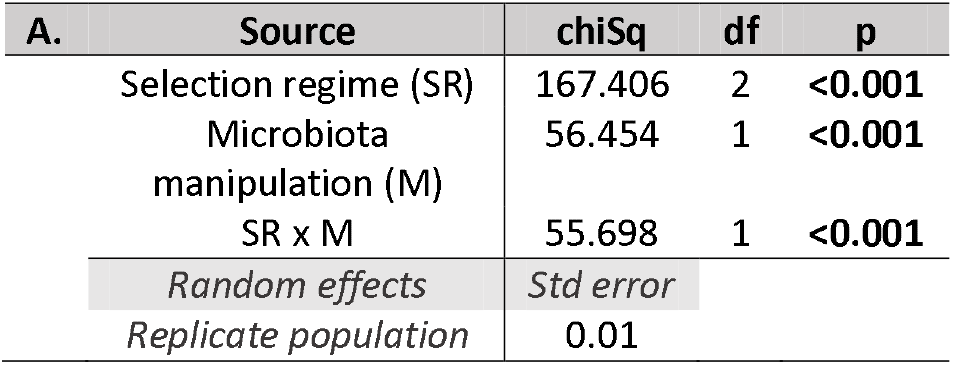
**A.**Summary of a generalized linear mixed effects model best fitted to Quasi-Poisson distribution for changes in the reproductive output across selection regimes (C, I and PI) as a function of microbiota manipulation (i.e., Normal vs UV-irradiated wheat). We specified the model as: Reproductive output ~ Selection regime x Microbiota manipulation + (1| Replicate population), with ‘selection regime’ and ‘Microbiota manipulation’ as fixed effects and ‘replicate population’ a as random effect.

